# Dynamic mode decomposition of resting-state and task fMRI

**DOI:** 10.1101/431718

**Authors:** Jeremy Casorso, Xiaolu Kong, Wang Chi, Dimitri Van De Ville, B.T. Thomas Yeo, Raphaël Liégeois

## Abstract

Component analysis is a powerful tool to identify dominant patterns of interactions in multivariate datasets. In the context of fMRI data, methods such as principal component analysis or independent component analysis have been used to identify the brain networks shaping functional connectivity (FC). Importantly, these approaches are *static* in the sense that they ignore the temporal information contained in fMRI time series. Therefore, the corresponding components provide a static characterization of FC. Building upon recent findings suggesting that FC dynamics encode richer information about brain functional organization, we use a dynamic extension of component analysis to identify *dynamic modes* (DMs) of fMRI time series. We demonstrate the feasibility and relevance of this approach using resting-state and motor-task fMRI data of 730 healthy subjects of the Human Connectome Project (HCP). In resting-state, dominant DMs have strong resemblance with classical resting-state networks, with an additional temporal characterization of the networks in terms of oscillatory periods and damping times. In motor-task conditions, dominant DMs reveal interactions between several brain areas, including but not limited to the posterior parietal cortex and primary motor areas, that are not found with classical activation maps. Finally, we identify two canonical components linking the temporal properties of the resting-state DMs with 158 behavioral and demographic HCP measures. Altogether, these findings illustrate the benefits of the proposed dynamic component analysis framework, making it a promising tool to characterize the spatio-temporal organization of brain activity.

## Introduction

The human brain exhibits a spatio-temporal organization of activity when performing a task (Rissman et al., 2004; Jiang et al., 2004; Gazzaley et al., 2004) and during resting-state (Greicius et al., 2003; Damoiseaux et al., 2006; Smith et al., 2013b). Characterizing the nature of interactions between different brain regions can be done via functional connectivity (FC) analyses of fMRI time series (Friston, 2011). FC is classically estimated within frameworks that ignore the temporal information that might be present in fMRI time series. The corresponding measures are called *static* because they are averaged over the entire fMRI time series and they neglect the ordering of fMRI time points (Theiler et al., 1992; Liégeois et al., 2017). Static measures of FC include Pearson correlation (Biswal et al., 1995; Buckner et al., 2009; Zalesky et al., 2010; Power et al., 2011; Yeo et al., 2011; Margulies et al., 2016), partial correlation (Fransson and Marrelec, 2008; Ryali et al., 2012), or mutual information (Chai et al., 2009), computed over entire fMRI time series. In the past years, converging evidence has suggested the presence of information beyond static FC and new frameworks exploiting this additional information have been proposed (Hutchison et al., 2013a; Preti et al., 2017). For example, sliding window correlation approaches explore the time-varying nature of FC (Hutchison et al., 2013b; Leonardi et al., 2013; Allen et al., 2014; Wang et al., 2016). Another way to extend the static FC framework is to include information encoded in the temporal ordering of time series, or precedence information, that was shown to be of particular relevance to describe fMRI dynamics (Roebroeck et al., 2011; Karahanoğlu and Ville, 2017). Measures exploiting precedence information include those using temporal derivatives (Shine et al., 2015; Karahanoğlu and Van De Ville, 2015; Bolton et al., 2018) or autoregressive parameters (e.g., Rogers et al., 2010) of fMRI time series. The latter extension of the static framework leads to *dynamic* measures of FC and the former may also be referred to as dynamic FC, or *time-varying* FC to avoid confusion (Liégeois et al., 2017).

In order to extract the most neurologically relevant information from FC measures, different approaches have been proposed. Among them, component analysis frameworks such as principal component analysis (PCA) (Viviani et al., 2005; Zhong et al., 2009), independent component analysis (Calhoun et al., 2009; Varoquaux et al., 2010; Smith et al., 2012), sparse PCA (Ulfarsson and Solo, 2007; Eavani et al., 2015) or constrained PCA (Hirayama et al., 2016) have been applied to identify the main patterns of interactions in fMRI time series. The identified components are interpreted as the main networks shaping brain connectivity, such as the default mode network, the visual network, or the motor network (Van Den Heuvel and Pol, 2010; Moussa et al., 2012). It is important to note that in these frameworks, components are identified from a static representation of the data. For example, principal components are the eigenvectors of the correlation -or covariance-matrix of the entire fMRI time series and can therefore be considered as static (Jolliffe, 1986). Based on this, and in the same way new FC measures were proposed to explore the time-varying nature of FC as well as its dynamic properties (Hutchison et al., 2013a; Preti et al., 2017), one could investigate the time-varying and the dynamic extensions of component analysis. The former can be achieved by applying a classical component analysis within a sliding window setting (Kiviniemi et al., 2011; Leonardi et al., 2013). The latter consists in exploiting the links between successive time points of time series in order to identify the main oscillatory modes driving their dynamics. Several methods, originally developed to characterize the spatio-temporal organization of climate systems, have been proposed to this end: principal oscillatory patterns (Penland, 1989; Bürger, 1993; von Storch et al., 1995), principal interaction patterns (Hasselmann, 1988), or more recently dynamic mode decomposition (Schmid, 2010). These methods are based on a first-order autoregressive (AR-1) representation of time series, and *dynamic modes* (DMs) are obtained by decomposing the AR-1 model matrix identified from the time series (see *Methods*). Since they are computed from a dynamic generative model, DMs not only have a spatial characterization, as in classical component analysis, but they also have a temporal characterization. More precisely, each DM is associated with a damping time and a period that provide further information about the dynamics of the main patterns of connectivity present in the data (Neumaier and Schneider, 2001).

The use of AR models to explore the multivariate interactions within fMRI time series dates back to more than a decade (Harrison et al., 2003; Valdés-Sosa, 2004; Valdes-Sosa et al., 2005; Rogers et al., 2010). The optimal order of these models decreases with the number of ROIs, and was in general found to be one in whole brain analyses considering more than a hundred ROIs (Valdes-Sosa, 2004; Ting et al., 2015). More recently, the AR-1 model was also shown to be a promising representation of FC dynamics (Liégeois et al., 2017). Therefore, we propose to explore the spatio-temporal organization of fMRI data using a dynamic extension of component analysis based on the AR-1 model of fMRI time series (Neumaier and Schneider, 2001). We start by illustrating the benefits of this approach over classical static component analysis on a toy example. Then, using resting-state and motor-task fMRI data of 730 healthy subjects of the Human Connectome Project (Van Essen et al., 2013), we compute the dominant DMs in resting-state and motor-task fMRI time series. Finally, using canonical correlation analysis (CCA), we explore the link between 158 HCP behavioral and demographic measures and the temporal properties of the resting-state DMs. Overall, our results complement previous findings using classical static methods, offering a new way to explore the spatio-temporal organization of brain function in rest and during task.

## Methods

### Data

We used data of the HCP 1200-subjects release comprising resting-state functional MRI, task functional MRI, and behavioral measures of young (ages 22–35) and healthy participants drawn from a population of siblings (Van Essen et al., 2013). All imaging data were acquired on a 3-T Siemens Skyra scanner using a multi-band sequence. Functional images have a temporal resolution of 0.72 *sec* and a 2-mm isotropic spatial resolution. For each subject, four 14.4 min runs (1200 frames) of functional time series were acquired (Smith et al., 2013a). Resting-state fMRI data was projected to the fs_LR surface space using the multimodal surface matching method (MSM-All; Robinson et al., 2013; Van Essen et al., 2013). Data were cleaned using the ICA-FIX method (Salimi-Khorshidi et al., 2014; Griffanti et al., 2014) and saved in CIFTI grayordinate format. Linear trends and mean cortical grayordinate signal were regressed. fMRI time series were parcellated into *N_R_* = 400 cortical regions of interest (ROIs) (Schaefer et al., 2017), demeaned and normalized ROI-wise. Finally, for consistency reasons we selected the *N_S_* = 730 subjects with four complete runs.

The HCP experimental protocol for the motor task was adapted from Buckner and colleagues (Buckner et al., 2011; Yeo et al., 2011). The motor-task fMRI data consisted of two 3.5 min (284 frames) runs per subject. Participants were presented with a visual cue, prompting them to perform one of the five motor tasks: squeeze left or right toes, tap left or right fingers, move tongue. Each task was performed twice within each run, with task block durations of 12 seconds preceded by a 3 second cue. We applied the same preprocessing as for resting-state fMRI time series, including a parcellation into 400 cortical areas (Schaefer et al., 2017), ROI-wise demeaning and normalization. The dynamic modes specific to each motor task were computed from sections of the time series: we selected only the last 6 seconds of each task block in order to consider the portion of the block where the hemodynamic response for the cued task is maximal, while also allowing a refractory period for the hemodynamic response of the previous task block (Buxton et al., 2004). Our results were reproduced for other choices of the fMRI time series subsections selected for each task (Figure S3).

Among the set of 458 HCP subject measures (SMs), we selected the 158 behavioral and demographic measures of most interest following Smith and colleagues (Smith et al., 2015). The list of SMs is reported in Table S1.

### Estimation of Dynamic Modes

We identify dynamic modes from the AR-1 representation of time series (Neumaier and Schneider, 2001):

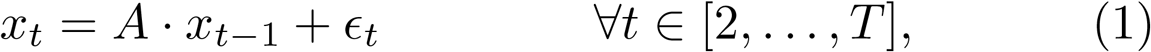

where 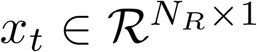 represents the fMRI time series at time *t*, 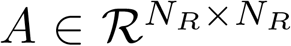 is the model parameter that encodes the linear relationship between successive time points, 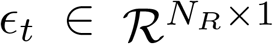 are the residuals of the model, and *T* is the number of time points. The model parameter *A* is computed by solving the following least-squares problem:

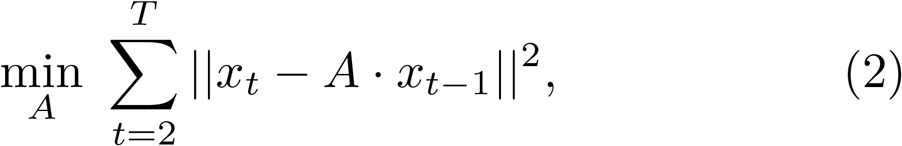

whose optimal solution is *A* = *XY*′(*YY*′)^−1^ where *X* = [*x*_2_,…,*x_T_*_−1_] and *Y* = [*x*_1_,…,*x_T_*_−1_] (Stoica and Moses, 2005). For subject-level computations, time series of different runs are concatenated and points corresponding to transitions between runs are removed from *X* and *Y*. The group-level estimation of *A*, denoted *A_G_*, is obtained by concatenating the time series of all runs and all subjects, and points corresponding to transitions between different runs or different subjects are removed from *X* and *Y*.

The eigendecomposition of *A* is defined as *A* = *S*Λ*S*^−1^ where the columns of *S*, denoted *S_i_*, are the eigenvectors of *A* and the diagonal matrix Λ encodes the corresponding eigenvalues, denoted *λ_i_*. Since *A* is not symmetric with real entries, *S* and Λ are in general complex with complex entries coming in complex conjugate pairs: if *λ_i_* is an eigenvalue and *S_i_* an eigenvector of *A*, then their complex conjugates are also eigenvalues and eigenvectors of *A*. Using this decomposition of *A*, one can reformulate the dynamic system (1) as a sum of linearly decoupled *dynamic modes* (DMs) (Neumaier and Schneider, 2001): each eigenvector of *A* defines one DM and the associated eigenvalue *λ_i_* provides its temporal characterization in terms of its damping time (Δ*_i_*) and period (*T_i_*):

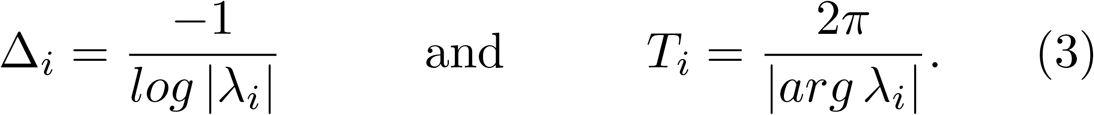

In summary, each DM is described by *S_i_* (spatial characterization), and Δ*_i_* and *T_i_* (temporal characterization). If *λ_i_* is complex or if it is real and negative, its period *T_i_* is bounded and the corresponding DM is known as an oscillator of period *T_i_*. If *λ_i_* is real and positive, its period *T_i_* is infinite and the associated DM is a relaxator characterized by a damping time Δ*_i_* (von Storch et al., 1995; Neumaier and Schneider, 2001). The modes are ordered by decreasing value of damping time Δ*_i_*, meaning that the least damped mode is assumed to be the most important one. For complex modes, we show both the real and imaginary parts of *S_i_* and do not show the corresponding redundant complex conjugate DM. In the same way real eigenvectors are defined up to a change of sign, complex eigenvectors are defined up to a random phase shift. To ensure unicity of complex eigenvectors *S_i_*, we impose orthogonality between their real and imaginary parts as detailed in Appendix A1 (see also Neumaier and Schneider, 2001). Values of Δ*_i_* and *T_i_* are multiplied by the temporal resolution of fMRI time series (0.72 *sec*) in order to express these quantities in *seconds*.

### Matching modes across subjects

In our last experiment we explore whether the temporal characterization of resting-state DMs encodes subject-specific behavioral and demographic information. However, subject-level DMs cannot be simply organized to allow comparison across subjects as there is no natural correspondence between DMs of different subjects. To circumvent this, we imposed the spatial modes (*S_i_*) of each subject to be equal to the group spatial modes. The temporal characteristics of these DMs in subject *k* are computed from the solutions of the following least-squares optimization problem:

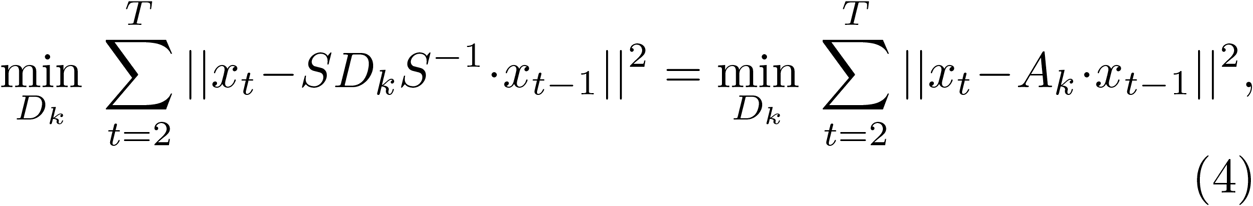

where *D_k_* is a diagonal matrix with entries to be optimized, *S* contains the eigenvectors of the optimal AR-1 parameter *A_G_*, and *A_k_* = *SD_k_S*^−1^. The difference between problem (4) and the unconstrained least-squares problem (2) is that *A_k_* is forced to share the same eigenvectors as A_G_. We provide the closed form expression of the solution to this problem in Appendix A2. From the optimal values of the entries of *D_i_* obtained from Eq. (A10)-(??), subject-specific damping times and periods of the DMs sharing the same spatial properties as the group DMs are computed using Eq. (3). In other words, for each subject we identify DMs that have the same spatial characterization as the group-level modes, but that have different damping times and periods. The dynamical properties of subject *i*’s resting-state DMs can be summarized by the damping times and periods of all its modes, resulting in a vector *v_k_* of length 2*N_R_* for each subject, where *N_R_* is the number of ROIs, equal to the number of DMs identified for each subject.

### Linking dynamic modes to behavior

We explore the link between the dynamical properties of resting-state DMs and the 158 HCP subject measures (SMs) using canonical correlation analysis (CCA) (Hotelling, 1936). The dynamical properties of DMs are encoded in the matrix 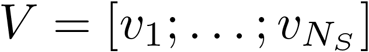 of size 2*N_R_* × *N_S_*, where *N_S_* is the number of subjects, and SMs are encoded in the matrix *B* of size 158 × *N_S_*. Age, gender and education were regressed from the SMs, and since periods *T_i_* may take infinite values we use the corresponding frequencies *f_i_* = 1/*T_i_* as inputs to CCA. In order to avoid CCA overfitting, we first applied PCA to both *V* and *B*, extracting the 36 first principal components (PCs) of *B* and the 20 first PCs of *V*, thereby keeping the same proportion of variance in both datasets: 97%. Our results are not significantly affected by changes in these values. The resulting 20 × *N_S_* and 36 × *N_S_* matrices were the inputs to the CCA framework. We evaluate the statistical significance of the identified canonical modes by permutation testing that preserves the family structure of the HCP dataset using the `hcp2blocks` function of Winkler and colleagues (Winkler et al., 2015).

The different analyses carried in this work are summarized in Figure 1.

**Figure 1:**
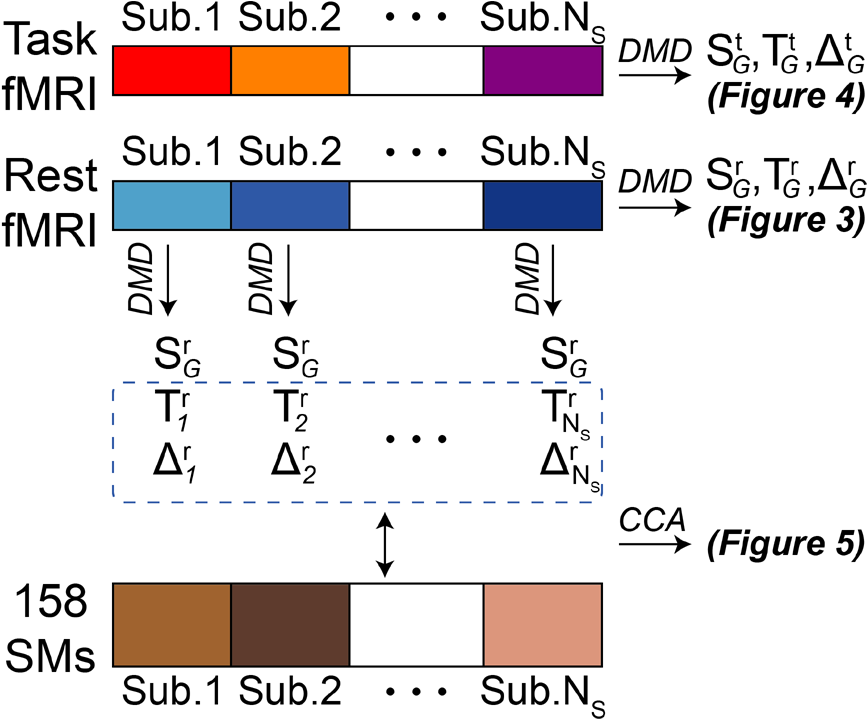
Summary of analyses carried in this work. Figure 3 and Figure 4 present the dominant group-level DMs in resting-state and in task, and Figure 5 presents the results of the CCA analysis linking 158 behavioral and demographic HCP measures to the temporal properties of the subject-level resting-state DMs. *CCA:* Canonical Component Analysis. *DMD*: Dynamic Mode Decomposition. *SMs*: Subject Measures.

## Results

### Dynamic modes vs. static components

In Figure 2 we present a toy example illustrating the spatial and temporal properties of the dynamic modes (DMs) of multivariate time series.

The toy example is composed of 5 variables -or regions of interest (ROIs)-containing two networks associated to different periods of oscillation. The first network, depicted in red in Figure 2A, oscillates with a period of 10 arbitrary units and is present in ROIs 1, 2 and 3 with ROIs 2 and 3 being dephased by π/7 with respect to ROI 1. The other network, depicted in blue in Figure 2A, oscillates with a period of 7 a.u. and is present in ROIs 3 and 4, with a dephasing of π/4. The networks overlap in ROI 3 which hence consists of a sum of two noisy sine waves with periods 7 and 10, and ROI 5 is pure white gaussian noise. The details of the simulation are found in Eq. (S1).

From these toy time series, we compute components -or modes-using three methods: two classical static component analysis frameworks (Figure 2B) and the dynamic extension that is used in this work (Figure 2C). Principal Component Analysis (PCA) and Independent Component Analysis (ICA) identify five components that recover more or less accurately the spatial organization of the two networks. In comparison, the proposed framework extracts five DMs including two pairs of complex conjugate DMs, thereby leading to the three unique DMs represented in Figure 2C. We first note that the spatial information recovered by the real parts of the DMs almost perfectly matches the organization of the two overlapping toy-networks. Then, the imaginary part of the DMs allows to recover the correct dephasing between the variables of a same DM from the ratio between its imaginary and real parts (Neumaier and Schneider, 2001): ROIs 2 and 3 of DM 1 are found to have a dephasing of π/7 with respect to ROI 1 of the same mode, and ROI 4 of DM 2 a dephasing of π/4 with respect to ROI 3 of the same mode, as modeled in the toy example (S1). Finally, the temporal properties of the DMs encode information about their characteristic damping times and periods. The oscillatory periods of the first and second DMs are 9.97 a.u. and 7.00 a.u., which almost exactly correspond to the true periods characterizing the two toy-networks. The third mode has an infinite period, which is consistent with the fact that white gaussian noise is not dominated by periodic oscillations. It also has a low damping time suggesting a weak influence between successive time points, as expected. On the contrary, the damping times of the first two modes are relatively high since the oscillatory behavior is not damped along the toy time series (Neumaier and Schneider, 2001). We finally note that the generative model of this toy example (Eq. (S1)) consists of sine functions which are *not* AR-1 time series as they contain information beyond the first autocorrelation lag. Hence recovering the correct toy-networks using the proposed dynamic component analysis framework does not result from considering a toy example fitting the framework’s assumptions, but rather highlights the relevance of exploiting the temporal information encoded in time series.

Overall, Figure 2 shows the additional information carried by dynamic modes as compared to classical static components such as independent or principal components. First, the complex-valued DMs allow to encode phase shifting information. Then, the associated eigenvalues provide a temporal characterization of the modes in terms of damping times and periods. These properties are not captured by classical components as their generative model treats time series as a collection of observations rather than as a succession of time points with a temporal structure (Jolliffe, 1986; Liégeois et al., 2017).

### Dynamic modes in resting-state fMRI

We compute group-level dynamic modes using resting-state fMRI data from 730 HCP subjects. The three dominant DMs, i.e. the ones with the longest damping times, are shown in Figure 3. In each DM, ROIs associated with warm colors are anti-correlated with those associated with cool colors.

The first mode displays two anti-correlated networks. The warm-colored network overlaps with the default mode network (DMN), including strong activations of the posterior cingulate cortex, the precuneus, and the ventromedial prefrontal cortex, as well as parts of the superior temporal sulcus and the inferior parietal lobule. The network associated with cool colors overlaps with the task positive network (TPN) with activations around the intraparietal sulcus, in the inferior temporal gyrus, and in lateral parts of the prefrontal cortex (Fox et al., 2005; Fransson, 2005; Yeo et al., 2011). This DM is found to be the one dominating resting-state fMRI dynamics, with a characteristic damping time of 7.27 *sec*. The second DM shows activations in lateral subregions of the DMN as well as in parts of the prefrontal cortex that were less or not activated in the first DM. These ROIs are anti-correlated with the primary visual area and the damping time of the DM is 5.87 *sec.* The third DM is complex-valued. The real part presents clear activations of the motor, somatosensory, and primary visual areas, with anticorrelation to regions of the lateral prefrontal cortex and the intraparietal sulcus. The imaginary part of this DM indicates a dephasing of brain activity in the orbitofrontal prefrontal cortex with respect to the areas activated in its real part. Subsequent DMs are shown in Figure S1.

**Figure 2:**
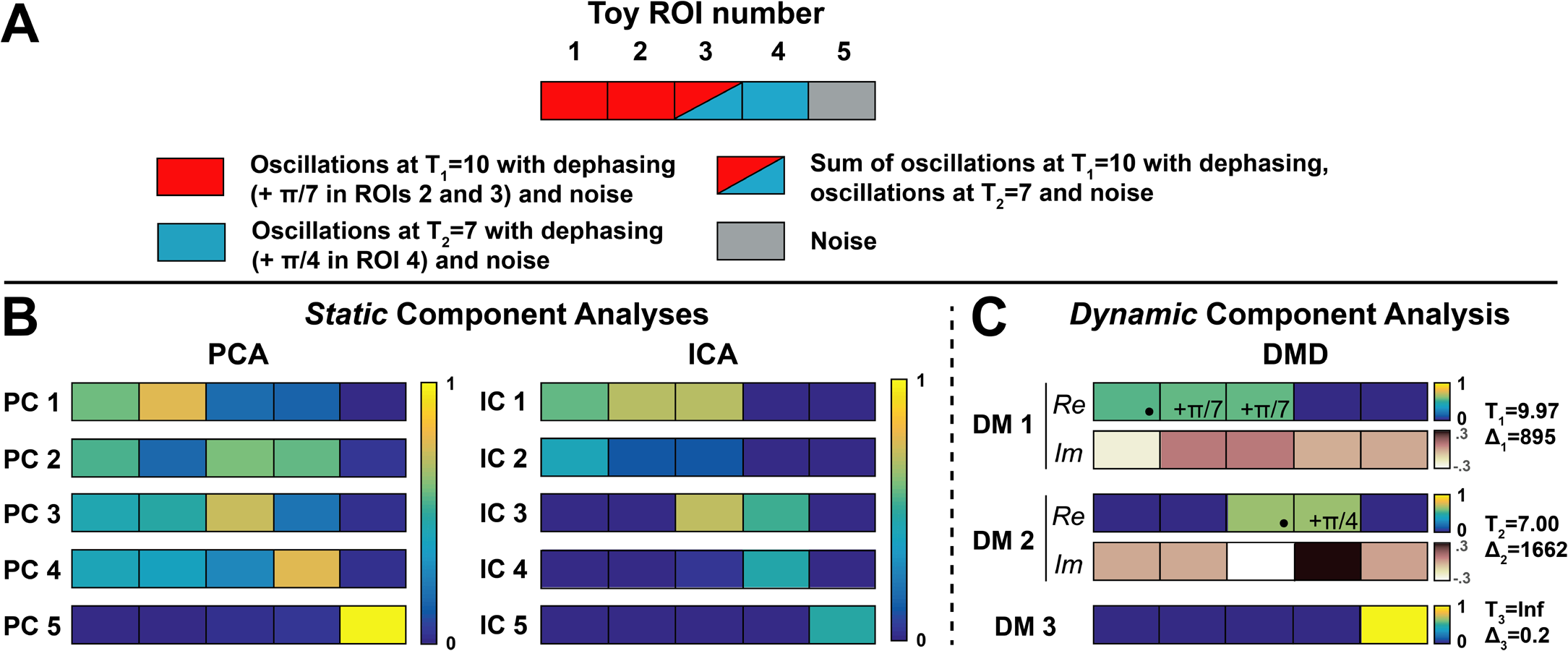
Toy example illustrating the differences between classical component analysis frameworks and their dynamic extension used in the present work. *(A)* The toy example consists of 5 variables representing activity in regions of interest (ROIs) as a function of time. ROIs are grouped into two overlapping networks based on their characteristic periods. The first network (red) is present in variables 1–3 and has a period of 10 (arbitrary units), and the second network (blue) is present in variables 3–4 with a period of 7 (a.u.). We included dephasing within networks, ROIs 2 and 3 being dephased by ð/7 as compared to ROI 1 within the red network, and ROI 4 being dephased by ð/4 as compared to ROI 3 within the blue network. ROI 5 is white gaussian noise. Details of the toy model are given in Eq. (S1). *(B)* Components identified using Principal Component Analysis (PCA) and Independent Component Analysis (ICA) on the toy time series. *(C)* A dynamic extension of component analysis identifies complex dynamic modes (DMs) that have a spatial and a temporal characterization. Only the first two DMs are complex, we represent their real *(Re)* and imaginary *(Im)* parts separately. The real parts of the DMs accurately recover the two original networks and their imaginary parts allow to recover dephasing within the networks. The DMs also have a temporal characterization in terms of periods *(T)* that are very close to the original values, and damping times (Ä).

**Figure 3:**
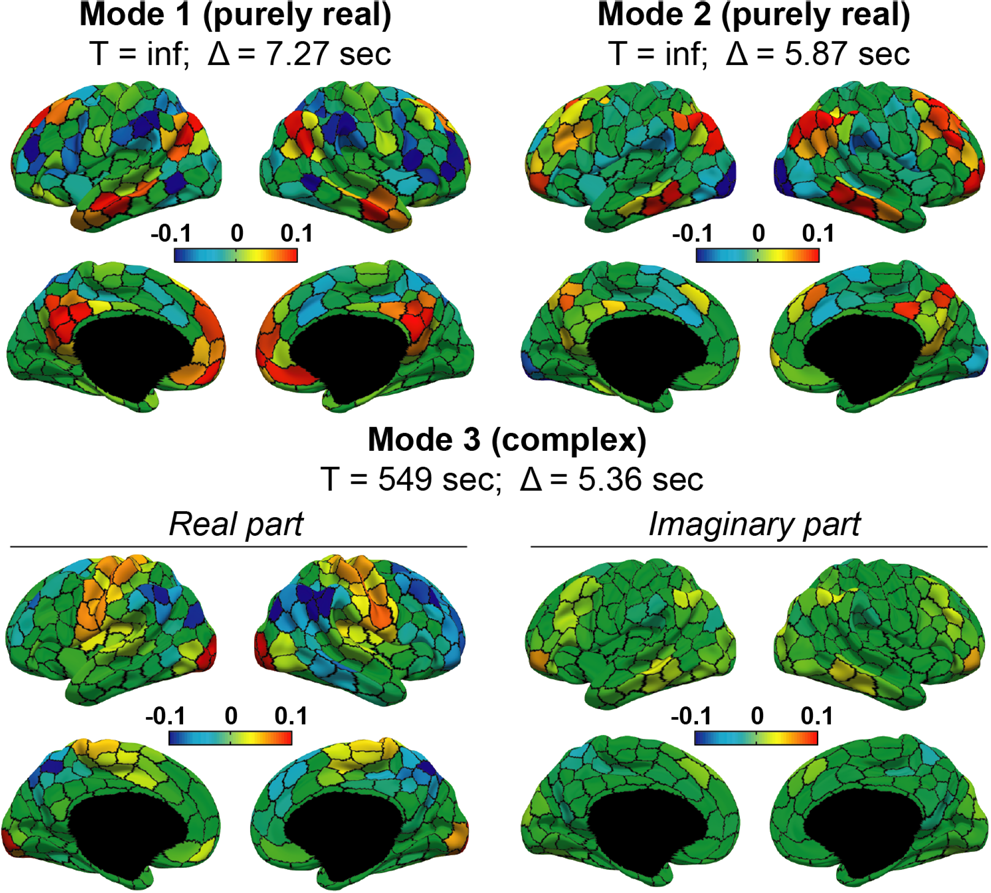
Dominant group-level resting-state dynamic modes. The first two modes are purely real and the third mode is complex. Modes oscillate with a characteristic period *T* and decay with a characteristic damping time Δ. The weight of each ROI in the dynamic modes is indicated by the colorbar, cool colors ROIs being anti-correlated with warm colors ROIs.

### Dynamic modes in task fMRI

The dominant DMs of fMRI time series acquired during five different motor task experiments are shown in Figure 4.

The five tasks involve moving left and right feet, left and right hands, and the tongue. The real parts of the dominant modes are consistent across tasks with the activation of the visual areas and the medial part of the posterior parietal cortex. Activations corresponding to classical activation maps of the tasks (Yeo et al., 2011; Barch et al., 2013) are encoded in the imaginary parts of the DMs along the motor and somatosensory cortex. This suggests the presence of a dephasing between specific task-related activity and the less specific activity observed in the real parts of the five dominant DMs. The identification of the task-related activity in the imaginary parts of the dominant DMs is robust to changes of parameters, as illustrated in Figure S3. The temporal properties of the dominant DMs are similar in the five experiments, with a characteristic damping time around 2.5 *sec* and an oscillatory period on the order of 12 *sec.* The second DMs in each task are presented in Figure S2.

**Figure 4:**
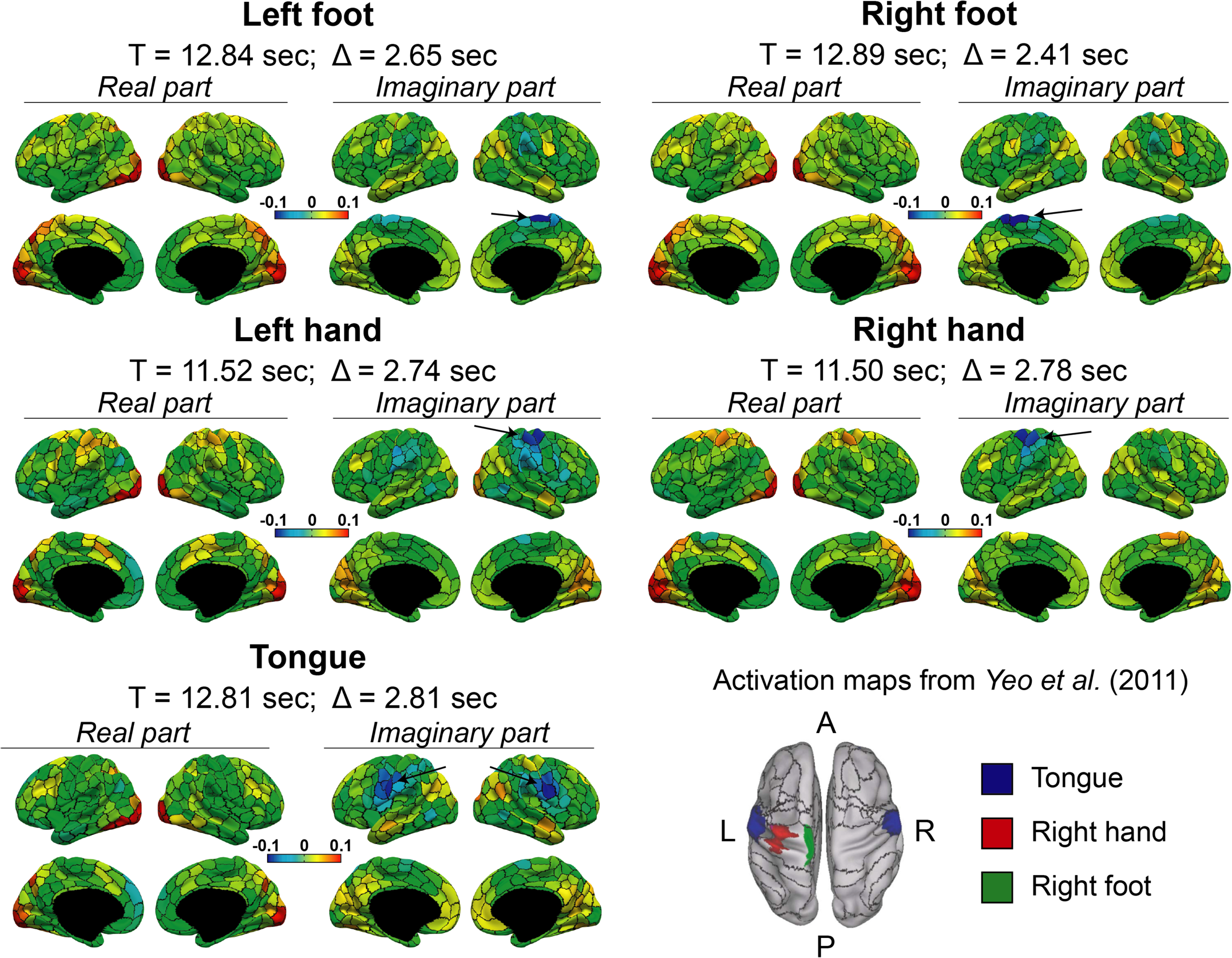
Dominant group-level dynamic modes in five motor-tasks: left foot, right foot, left hand, right hand, and tongue. The dominant mode is complex in each task, and the real and imaginary parts are displayed separately on the cortical surface map. Classical activation maps corresponding to these tasks are represented in the bottom right (adapted from Yeo et al. (2011), with permission). Task-related activity classically identified in activation maps seems to be mostly encoded in the imaginary parts of the DMs (arrows), indicating a dephasing of activity with respect to the corresponding real parts.

### Linking resting-state DM’s temporal characteristics to behavior

In order to further explore the information carried by the temporal properties of DMs, we perform a canonical correlation analysis (CCA) between, on the one hand, the subject-specific loadings -from which temporal characteristics are computed, see *Methods* - of group-level spatial modes, and on the other hand, 158 HCP behavioral and demographic subject measures (SMs).

**Figure 5:**
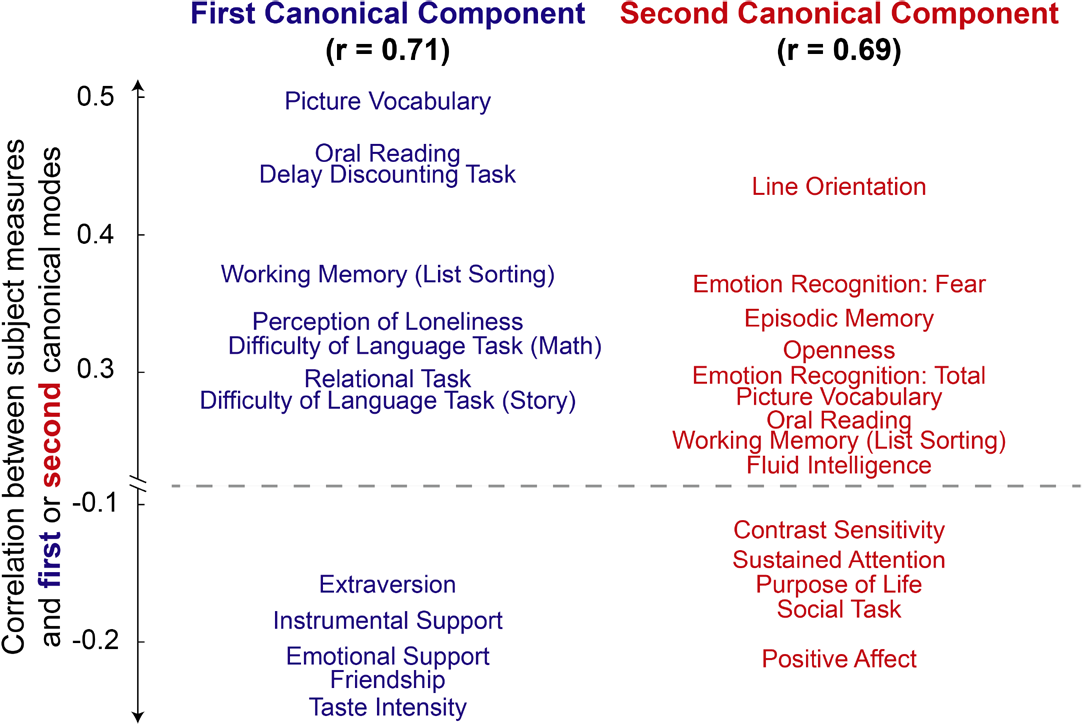
Most positive and negative correlations between the subject measures (SMs) and the two canonical modes linking 158 behavioral measures and the temporal characteristics of resting-state DMs. Full HCP headers corresponding to the SMs are found in Table S1

We find two statistically significant (*p <* 0.05) canonical modes linking SMs and temporal characteristics of resting-state DMs with correlations of *r* = 0.71 and *r* = 0.69. SMs that most positively or negatively correlate with the two canonical modes are shown in Figure 5. The exact values of the correlations with the CCA modes as well as the full HCP denominations are reported in Table S1.

## Discussion

In this study, we propose a data-driven approach to identify dynamic modes (DMs) in rest and task fMRI time series. In the same way new methods have been recently developed to exploit the dynamic nature of FC time series (Hutchison et al., 2013a; Preti et al., 2017), the proposed framework can be seen as a dynamic extension of classical component analysis, allowing to identify dynamic brain networks -or dynamic modes-that offer a richer characterization of the main patterns shaping brain activity.

### Related methodological approaches

Various component analysis methods such as principal component analysis (PCA) or independent component analysis (ICA) have been applied to fMRI time series in order to identify the main interaction patterns driving functional connectivity (FC) (Viviani et al., 2005; Ulfarsson and Solo, 2007; Calhoun et al., 2009; Smith et al., 2012; Eavani et al., 2015; Hirayama et al., 2016). Importantly, these component analysis frameworks are *static* in the sense that they treat successive time points of multivariate time series as independent observations. For example, PCA relies on the eigendecomposition of the correlation matrix of time series which is a static measure of FC (Jolliffe, 1986; Theiler et al., 1992) and ICA uses static measures such as kurtosis to maximize non-gaussianity of independent components (Hyvarinen et al., 2001). In contrast, the proposed framework is based on a first-order multivariate autoregressive (AR-1) model of fMRI time series which is *dynamic* because it exploits the statistical link between successive time points (Theiler et al., 1992). Building upon this key property, multivariate AR-1 models were shown to capture much more dynamic FC information as compared to their static counterparts on which classical components frameworks are based (Liégeois et al., 2017).

Our approach should also be distinguished from the time-resolved frameworks consisting in performing classical component analysis within sliding windows (Kiviniemi et al., 2011; Leonardi et al., 2013). Indeed, these methods exploit the sample variability of static measures along time series to derive the temporal fluctuations of these measures; i.e., within a window only static measures of connectivity are used. On the contrary, our framework does not provide time-resolved information and instead exploits the whole time series in order to provide the most accurate estimates of the *dynamical* connectivity patterns shaping the time series. This distinction is reminiscent of the distinction between time-varying and dynamic measures of FC mentioned in the *Introduction* and detailed in previous work (Liégeois et al., 2017). For the same reason, activation time series that can be associated to principal or independent components (e.g., Calhoun et al., 2009) are essentially different from the proposed framework as they carry time-resolved information of *static* components.

As for principal components, DMs are most often ordered by decreasing absolute value of the associated eigenvalues. In the context of PCA, the eigenvalue carries information about the amount of variance of the original data explained by a component (Jolliffe, 1986). In the case of DMs, the absolute value of the associated eigenvalues finds an interpretation in terms of damping time, which is why the DMs with highest absolute eigenvalue; i.e., the ones with longest damping times, are ranked first. The distribution of damping times of the first 100 DMs Figure is shown in S4. Note that other ordering rules exploiting the amplitude of the noise in the AR-1 model (1) have been proposed (Neumaier and Schneider, 2001).

### Dynamic modes in rest and task

The dominant resting-state DMs presented in Figures 3 and S1 all have connections with known resting-state networks. The first resting-state DM contains two anticorrelated networks: the default mode network (DMN - warm colors in Figure 3) and a set of ROIs (cool colors) overlapping, but not exactly matching, with the task positive network (TPN) defined by Fox and colleagues (Fox et al., 2005). Identifying the DMN in the least damped DM might contribute to explain its robust identification in resting-state fMRI data (Van Den Heuvel and Pol, 2010; Moussa et al., 2012), as well as its central role in the macroscale cortical organization (Margulies et al., 2016; Gu et al., 2017). The second DM shows strong activations in lateral parts of the DMN that were not activated in the first DM, with anti-correlation localized around the visual network. This suggests that DMs 1 and 2 capture different sub-networks of the DMN and might shed a new light into the complex organization of the DMN and its interactions with other networks (Uddin et al., 2009; Raichle, 2015; Karahanoğlu and Van De Ville, 2015). The next DMs show activations in the motor and somatosensory areas around the central sulcus (DMs 3 and 4), in the visual area (DMs 4 and 5) and in subregions of the DMN (DM 5), thereby also providing a characterization of the interactions between known resting-state networks. Finding neurologically interpretable dominant DMs suggests that the proposed decomposition and the corresponding ranking of DMs based on their damping times exploit important features of fMRI time series. In comparison, independent components of fMRI time series cannot be ranked and need to be classified as being of neurological origin or not (Tohka et al., 2008; Beckmann, 2012; Griffanti et al., 2017).

Atop of the spatial properties of resting-state DMs, each mode is also associated with a damping time and a period. This temporal characterization of the modes is not directly available for static components as they are not computed from a dynamic model of fMRI time series. The utility of the spatial properties of resting-state DMs has been illustrated in Figures 3 and 4. We further discuss in a later subsection the use of their associated temporal properties by linking them to demographic and behavioral subject measures.

In task condition, the dominant DM is complex and shows activations in the motor and somatosensory cortex that are dephased with the visual area and the posterior parietal cortex (Figure 4). The DMN is partially found in the second task DM and seems decoupled from its anticorrelated network found in the dominant resting-state DM. Unlike in the resting-state, the DMN is not the dominant DM which is consistent with the known decreased activity of the DMN during externally-oriented tasks (Fox et al., 2005). Interestingly, some of the regions that were anti-correlated with the DMN in resting-state DMs, such as the primary visual area, are consistently activated in task DMs which suggests they might play a key role in the transition between rest and task conditions.

### Dephasing as a fingerprint of task-related activity

The activation maps of the five motor tasks considered in this work have been studied previously, showing activations along the primary motor and somatosensory cortex (Yeo et al., 2011; Barch et al., 2013). The same ROIs are found to be active in the imaginary parts of the dominant task DMs, together with activations in the visual cortex and in the posterior parietal cortex encoded in the real parts of the DMs (Figure 4). The imaginary and real parts of a DM represent aspects of the mode that share the same damping times and periods, but that are dephased by π/2 (von Storch and Zwiers, 1999; Neumaier and Schneider, 2001). Activations of the visual and posterior parietal cortices in the same DM as the somatomotor cortex suggest that these areas are also involved in the execution of the task. This might be explained by the known role of the posterior parietal cortex as a sensorimotor interface transforming visual information into motor commands (Andersen and Buneo, 2002; van Mier et al., 2004; Buneo and Andersen, 2006; Dean et al., 2012). The dephasing between the active areas of the real and imaginary parts of the dominant DM could be due to latency differences of activations in these ROIs, as previously found in a visuomotor task (Lin et al., 2013). A change in the hemodynamic response function could also contribute to this dephasing (Bellgowan et al., 2003; Handwerker et al., 2012; Orban et al., 2015; Goelman et al., 2017) and delineating its exact causes would require further examination, for example using multi-modal data (Lewis et al., 2016).

Most studies use a general linear model to identify task-related activations (Yeo et al., 2011; De Guio et al., 2012; Barch et al., 2013; Kristo et al., 2014). In the case of motor tasks such as the ones considered here, this amounts to identifying task-related activity by comparing brain activity at rest and during task. In contrast, the DMs presented in Figures 4 and S3 are computed only from the time series acquired when performing the task, and no comparison with a baseline condition is used. Recovering the classical activation maps of these tasks in the dominant DMs therefore suggests that the proposed framework -and the associate ranking criterion of the DMs-exploits relevant features of fMRI time series. This could also explain why ROIs that are less directly related to the task execution, and are therefore not found in classical activation maps, are found to be active in the dominant DMs. Overall, dominant DMs showing strong dephasing across brain regions could be another element distinguishing task from resting-state fMRI time series (Zhang et al., 2016), possibly contributing to our understanding of the mechanisms associated with various task conditions.

### Behavioral counterparts of dynamic modes

The DMs have a *spatial* characterization, encoded in the eigenvectors of the matrix *A* in model (1), and the corresponding eigenvalues provide a *temporal* characterization of the DMs in terms of damping times and periods (Eq. (3)). The previous analyses mainly explored the spatial properties of the DMs. In Figure 5, we test whether the temporal characteristics of the DMs are linked to 158 HCP behavioral and demographic subject measures (SMs) using CCA, and identify two significant CCA modes. To this end we impose the subject-specific DMs to share the same eigenvectors, i.e. the same spatial properties, as the group DMs, only allowing for inter-subject variability of the corresponding temporal properties. This is quite restrictive as individual network topography was shown to be qualitatively different from group average estimates (Braga and Buckner, 2017; Gordon et al., 2017; Kong et al., 2018). However, we decided to specifically explore the interpretation of the *temporal* properties of the DMs in order to complement our previous findings. A combined analysis linking SMs to both spatial and temporal properties of DMs is left for future work, as detailed further.

We use the same SMs as Smith and colleagues (Smith et al., 2015) and follow a similar methodology. Smith and colleagues found one significant CCA mode relating a static measure of functional connectivity and the SMs, we find two (Figure 5). Interestingly, the three SMs most positively correlated with our first CCA mode (Picture Vocabulary Test, Delay Discounting and Oral Reading Recognition) are also strongly positively correlated with Smith and colleagues’ first CCA mode. Hence these SMs are related to both static and dynamic aspects of functional connectivity, which further evidences their links with brain functional organization. The other SMs most strongly correlating with our first CCA mode are different from Smith and colleagues’ and seem to draw a distinction between measures related to a classical intelligence factor (Spearman, 1904), which are positively correlated with the first CCA mode, and measures closer to life satisfaction, which are negatively correlated. The second CCA mode shows strongest correlations with SMs such as the number of correct identification of various emotions, Fluid Intelligence, Purpose of Life, Social Task Performance, and Positive Affect which evaluates the level of pleasurable engagement including happiness, joy, enthusiasm, and contentment (Van Essen et al., 2013). As such, this CCA mode could be related to the concept of emotional intelligence that refers to the cooperative combination of intelligence and emotion (Salovey and Mayer, 1990; Roberts et al., 2001; Mayer et al., 2008). This includes the ability to recognize emotions and use it to enhance thoughts. Emotional intelligence was shown to be linked to happiness (Furnham and Petrides, 2003), life satisfaction (Ciarrochi et al., 2000) and quality of social relationships (Lopes et al., 2004) which were all found to be strongly correlated to the second CCA mode. These results offer a new perspective on the behavioral information encoded in resting-state FC, complementing previous findings using classical (i.e., static) measures of brain functional organization.

### Methodological limitations and future directions

Adding a dynamic dimension to the classically static component analysis framework modifies the properties of the resulting components that need to be interpreted accordingly (Neumaier and Schneider, 2001). The features of DMs have been discussed along this study but some interpretations in this application could be further investigated. For example, Figure S3 shows the dominant DMs computed from a subsection of the fMRI time series used to compute the dominant DMs presented in Figure 4. The spatial distribution of the modes are very similar, with activations of the motor and somatosensory ROIs encoded in the imaginary parts of the DMs, but the damping times and periods show substantial differences in the two cases. In general, our experiments (not shown) suggest that the period of the dominant DM increases with the length of the time series used to perform the decomposition, sometimes leading to a purely real dominant DM with infinite period for long fMRI time series (e.g., when several runs are concatenated). As a consequence, inter-subjects comparisons of these quantities should be performed using time series of same length for each subject, as in our CCA analysis. This effect is puzzling and its causes are unclear. One possible explanation is that the frequencies of oscillating patterns are fluctuating along the fMRI time series (Chang and Glover, 2010; He, 2011) which might affect the correct identification of a single frequency to characterize the dominant DM in long fMRI time series. Pre-filtering fMRI time series in specific frequency bands might help to circumvent this limitation.

A time-resolved computation of the DMs using a sliding window approach could be considered but three limitations need to be emphasized. First, the interpretation of DMs is more complex than classical static components and the interpretation of the temporal fluctuations of DMs would be even more, possibly masking the important properties carried by DMs. Second, the identification of an AR-1 model of multivariate time series (Eq. (1)) requires at least as many time points as variables (Stoica and Moses, 2005). Using this study’s parcellation, this would result in windows containing at least 400 fMRI time points which is likely to be too long to capture meaningful temporal fluctuations of DMs (Liégeois et al., 2016; Preti et al., 2017). Note that low rank approximations of A, the model matrix of the AR-1 model, can be estimated using less time points, thereby identifying only the most important DMs (Neumaier and Schneider, 2001). This might be sufficient as only a small proportion of DMs have high damping times (Figure S4). Finally, we showed in previous work that a single AR-1 model of fMRI time series captures a significant part of FC temporal fluctuations (Liégeois et al., 2017), thereby suggesting that a time-resolved computation of DMs might not be necessary to capture the dominant spatio-temporal properties of functional connectivity.

We explore the link between the temporal properties of DMs, and demographic and behavioral SMs in order to illustrate their utility and complementarity with the spatial properties of DMs. Subsequent studies focusing on the link between resting-state dynamic FC and SMs could use both spatial and temporal properties of DMs in their analysis, and/or use alternative methods to CCA (Grellmann et al., 2015). Using only the top DMs might be another way to reduce the dimensionality of this multivariate analysis problem, but this depends on the ranking criterion of the DMs. Even if the criterion we use seems to be meaningful in this application, other rankings could be considered (Section 2.1, Neumaier and Schneider, 2001).

## Conclusion

This study uses a *dynamic* extension of component analysis to compute the main dynamic modes (DMs) shaping brain function in different conditions. Using data of 730 HCP subjects, we compute the dominant group DMs in resting-state and in five different motor-task conditions. Resting-state DMs show similarities with classical (i.e., *static*) resting-state networks, but also significant differences that provide further insight about their interactions and dynamical properties. The dominant task DMs show activity in the areas of the primary somatomotor cortex classically identified in activation maps of the five tasks. The same DMs also consistently show dephased activity in other ROIs including the visual and posterior parietal cortices, revealing the dynamic interactions between these brain areas during the execution of the tasks. Finally, a CCA analysis identified two CCA modes linking the temporal properties of DMs and 158 HCP SMs, thereby complementing previous findings that identified a single mode of co-variation between the same SMs and static markers of FC. Altogether, these results suggest that dynamic mode decomposition is a promising tool to explore the spatiotemporal properties of brain functional dynamics encoded in fMRI time series.

## Acknowledgements

This research was supported in parts by the CHISTERA IVAN project (20CH21 174081), the Center for Biomedical Imaging (CIBM) of the Geneva - Lausanne Universities and the EPFL, the Singapore MOE Tier 2 (MOE2014-T2–2–016), the NUS Strategic Research (DPRT/944/09/14), the NUS SOM Aspiration Fund (R1850002–71720), and the Neuroimaging Informatics and Analysis Center (1P30NS098577). Data were provided by the Human Connectome Project, WU-Minn Consortium (Principal Investigators: David Van Essen and Kamil Ugurbil; 1U54MH091657) funded by the 16 NIH Institutes and Centers that support the NIH Blueprint for Neuroscience Research; and by the McDonnell Center for Systems Neuroscience at Washington University.

## Appendix

### Al. Unicity of complex eigendecomposition

Let us denote 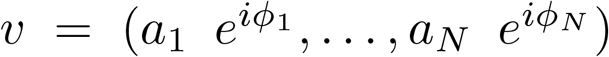 a complex eigenvector of a square matrix *A* of size *N*, with *a_i_* and *ϕ_i_* denoting the amplitude and phase of its *i^th^* entry. By definition, *v* has unit norm and we have *Av* = *λv*, where *λ* is the eigenvalue associated to *v*. Any vector *v_α_* = *v* · *e^iα^* also has a unit norm and verifies *Av_α_* = *λv_α_*. In order to have a unique representation of each eigenvector, we follow the convention of Neumaier and Schneider imposing that the real and imaginary parts of all eigenvectors are orthogonal, and that the amplitude of the imaginary part is smaller than or equal to the amplitude of the real part (Neumaier and Schneider, 2001). First, we identify the phase α_0_ that leads to orthogonality between the real and imaginary parts of 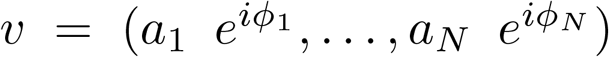:

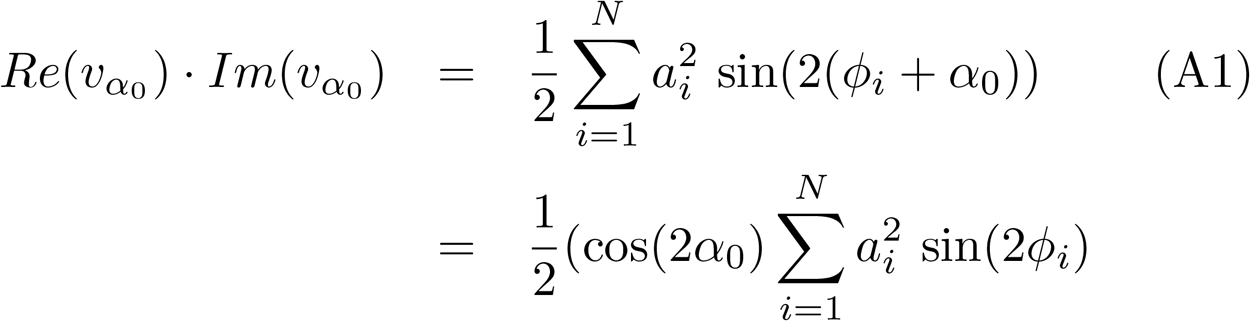

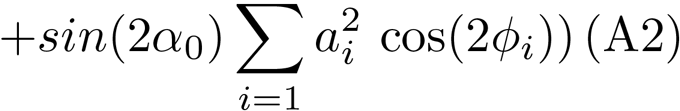

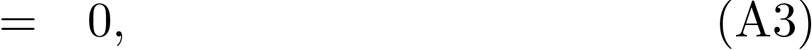

which leads to:

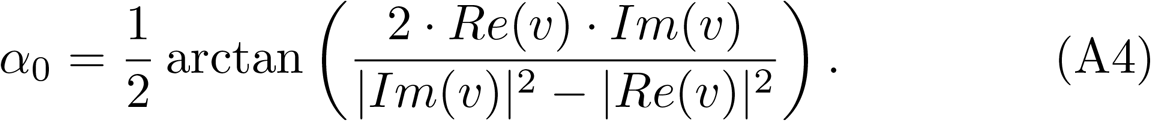

Since *α*_0_ is defined up to shifts of π/2, the real and imaginary parts of 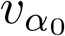 can be interchanged in order to make the absolute value of its real part larger or equal to its imaginary part.

### A2. Identification of spatially matched DMs

We solve the following optimization problem:

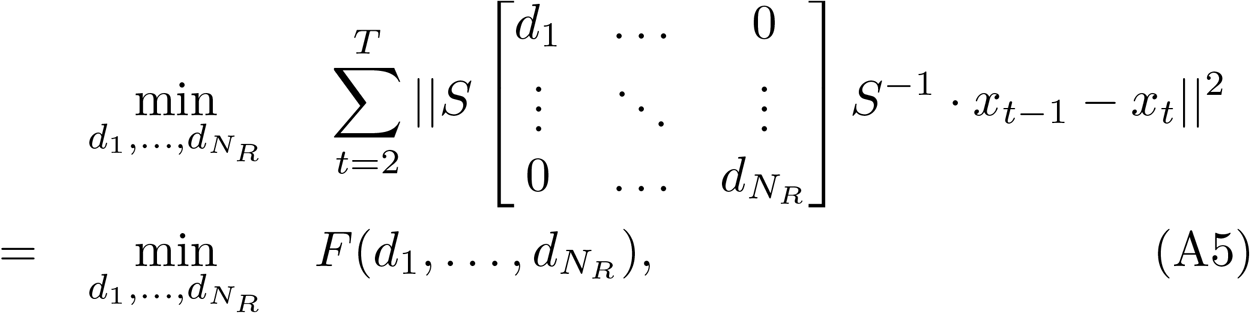

where *S* is a square matrix of size *N_R_*. Denoting *u_n_* (*v_n_*) the *n^th^* column (line) of *S* (*S*^−1^), we can write:

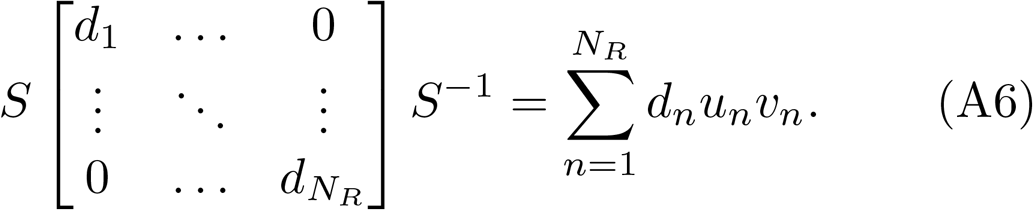

Using Eq. (A6), the argument of the minimization problem (A5) writes

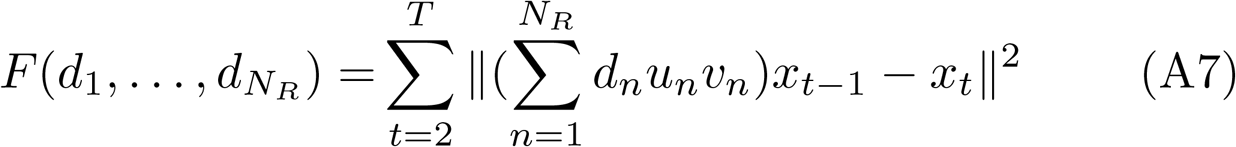

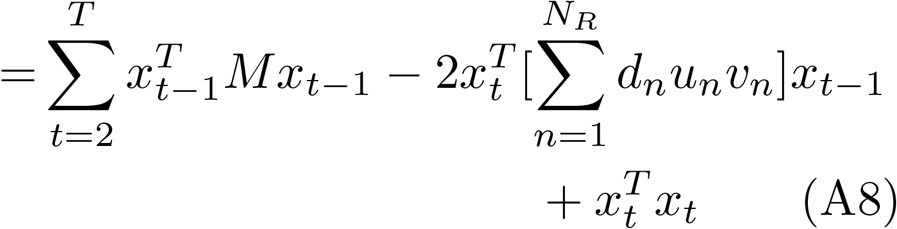

where *M* is given by:

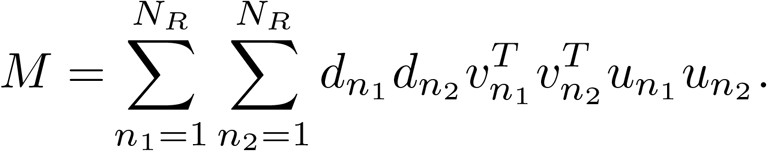

The solution of the convex problem (A5) is found by finding the zeros of its partial derivatives with respect to *d_i_,* ∀*i* ∈ {1,…,*N_R_*}:

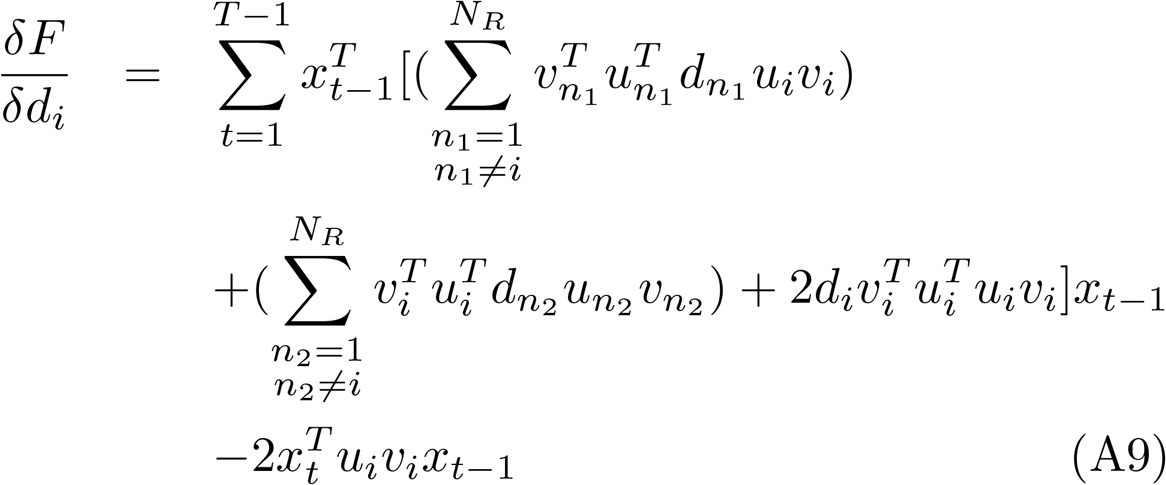

which leads to a system of *N_R_* linear equations providing the optimal value of the *N_R_* unknowns:

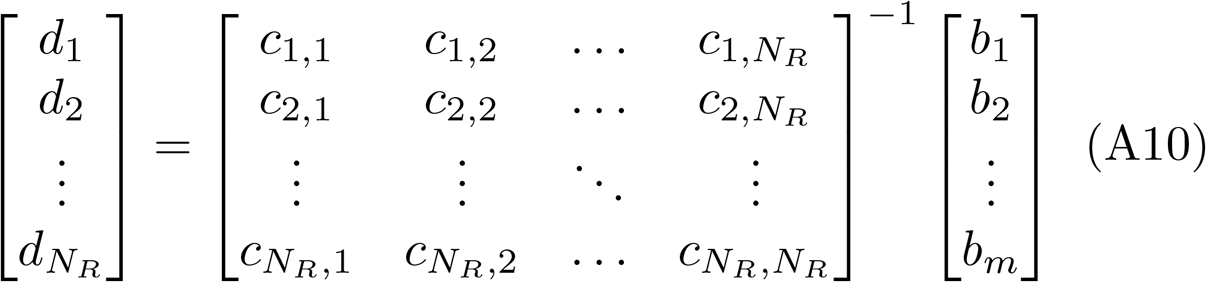

where 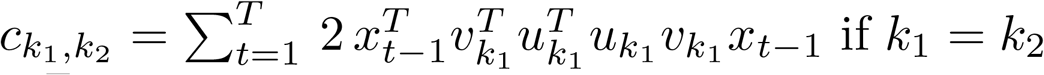, 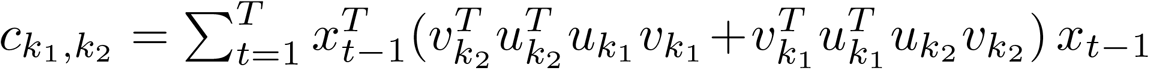 otherwise, and 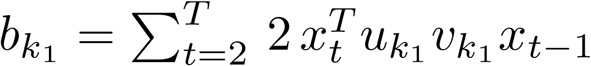.

## Supplementary Material

### Toy model

The generative model of the 5-variate time series used in Figure 2 is, for *t* ∈ [1,…, 1000]:

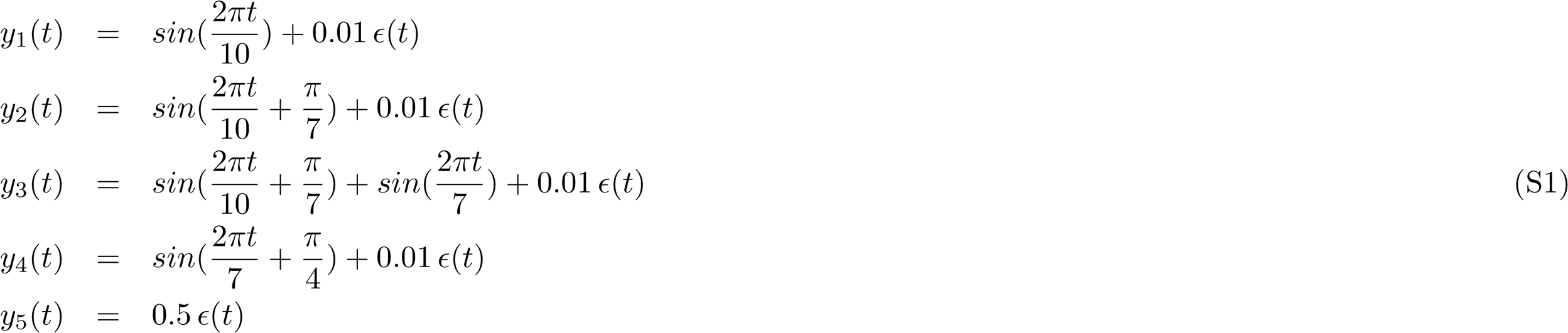

where *ε*(*t*) is white gaussian noise.

### Detail of subject measures

We use the 158 HCP subject measures that were previously used by Smith and colleagues (Smith et al., 2015): PicVocab_Unadj PicVocab_AgeAdj PMAT24_A_CR DDisc_AUC_200 THC LifeSatisLUnadj List Sort_AgeAdj Read-Eng_Unadj SCPT_SPEC ReadEng_AgeAdj ListSort_Unadj DDisc_AUC_40K Avg_Weekday_Any_Tobacco_7days Num_Days_Used_Any_Tobacco_7days Total_Any_Tobacco_7days PicSeq_AgeAdj FamHist_Fath_DrgAlc Pic-Seq_Unadj Avg_Weekday_Cigarettes_7days Avg_Weekend_Any_Tobacco_7days Total_Cigarettes_7days Dexterity_AgeAdj Avg_Weekend_Cigarettes_7days Dexterity_Unadj Times_Used_Any_Tobacco_Today PSQI_Score AngAggr_Unadj Taste_AgeAdj ASR_Rule_Raw Taste_Unadj ASR_Thot_Raw EVA_Denom SSAGA_TB_Still_Smoking FamHist_Fath_None ASR_Thot_Pct PercStress_Unadj ProcSpeed_AgeAdj ASR_Rule_Pct ProcSpeed_Unadj DSM_Antis_Raw ER40_CR NEOFAC_A ASR_Crit_Raw VSPLOT_TC NEOFAC_O ER40ANG VSPLOT_OFF SSAGA_Times_Used_Stimulants ASR_Soma_Pct SSAGA_Mj_Times_Used DSM_Antis_Pct Ca-rdSort_AgeAdj ASR_Extn_Raw ASR_Oth_Raw ASR_Totp_T ASR_Extn_T ASR_Totp_Raw EmotSupp_Unadj DSM_Anxi_Pct PercReject_Unadj ER40NOE DSM_Anxi_Raw ASR_TAO_Sum SSAGA_TB_Smoking_History CardSort_Unadj PosAffect_Unadj SSAGA_ChildhoodConduct Odor_AgeAdj ASR_Witd_Raw SSAGA_Alc_Hvy_Frq_Drk ASR_Soma_Raw DSM_Depr_Pct ASR_Aggr_Pct SSAGA_Alc_12_Max_Drinks DSM_Depr_Raw Mars_Final PercHostiLUnadj DSM_Somp_Pct SSAGA_Alc_Age_1st_Use ASR_Witd_Pct IWRD_TOT PainInterLTscore MMSE_Score SSAGA_Alc_12_Frq_Drk Odor_Unadj SSAGA_Alc_D4_Ab_Sx SSAGA_Mj_Use ASR_Aggr_Raw SSAGA_Mj_Ab_Dep DSM_Somp_Raw FearSomat_Unadj SSAGA_Alc_12_Drinks_Per_Day Mars_Log_Score Self-Eff_Unadj SCPT_SEN NEOFAC_N SSAGA_Agoraphobia ASRJntn_T AngHostil_Unadj Num_Days_Drank_7days SSAGA_Times_Used_Cocaine Loneliness_Unadj ASR_Intn_Raw SSAGA_Alc_Hvy_Drinks_Per_Day MeanPurp_Unadj DSM_Avoid_Pct NEOFAC_E Total_Beer_Wine_Cooler_7days DSM_Avoid_Raw Avg_Weekday_Wine_7days Flanker_AgeAdj ASR_Anxd_Pct Avg_Weekend_Beer_Wine_Cooler_7days SSAGA_Alc_D4_Ab_Dx Total_Drinks_7days SSAGA_Alc_Hvy_Max_Drinks FearAffect_Unadj Total_Wine_7days Avg_Weekday_Drinks_7days ER40SAD Flanker_Unadj ER40FEAR Avg_Weekday_Beer_Wine_Cooler_7days SSAGA_Times_Used_Illicits Avg_Weekend_Drinks_7days SSAGA_Alc_D4_Dp_Sx NEOFAC_C Total_Hard_Liquor_7days Correction SSAGA_Alc_Hvy_Frq_5plus DSM_Adh_Pct ASR_Attn_Pct VSPLOT_CRTE SSAGA_Depressive_Ep AngAffect_Unadj SSAGA_PanicDisorder Avg_Weekend_Hard_Liquor_7days FamHist_Moth_Dep ASR_Anxd_Raw SSAGA_Times_Used_Opia-tes SSAGA_Times_Used_Sedatives SSAGA_Alc_Hvy_Frq SSAGA_Alc_12_Frq_5plus Friendship_Una-dj SSAGA_Depressive_Sx ASR_Attn_Raw ASR_Intr_Raw SSAGA_Alc_12_Frq FamHist_Fath_Dep InstruSupp_Unadj ASRJntr_Pct SSAGA_Times_Used_Hallucinogens Avg_Weekend_Wine_7days Fa-mHist_Moth_None Sadness_Unadj DSM_Hype_Raw DSM_Adh_Raw DSMJnat_Raw

The details of subject measures presented in Figure 5 are reported in Table S1.

**Table S1:**
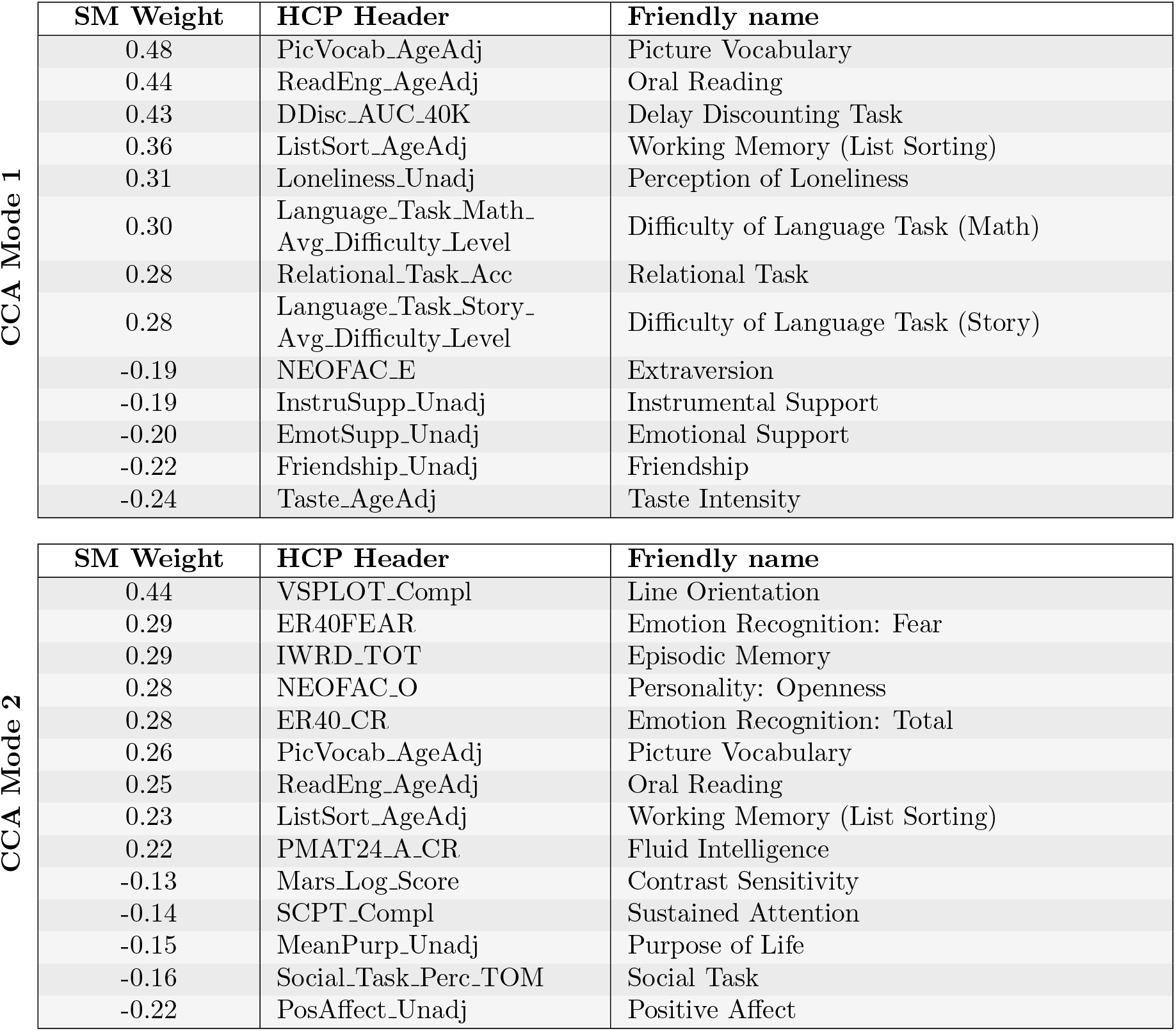
List of subject measures (SMs) that are most strongly associated with the two significant CCA modes. HCP headers and names used in Figure 5 are given, and more details about the SMs can be found at https://wiki.humanconnectome.org/display/PublicData/. The SM weights are determined by the correlation between the SMs and the two canonical modes linking the 158 SMs and the temporal characteristics of resting-state DMs.

### Additional modes found in resting-state and task conditions

Figure S1 shows the fourth and fifth group-level dynamic modes of resting-state fMRI time series. The first three modes are shown in Figure 3.

**Figure S1:**
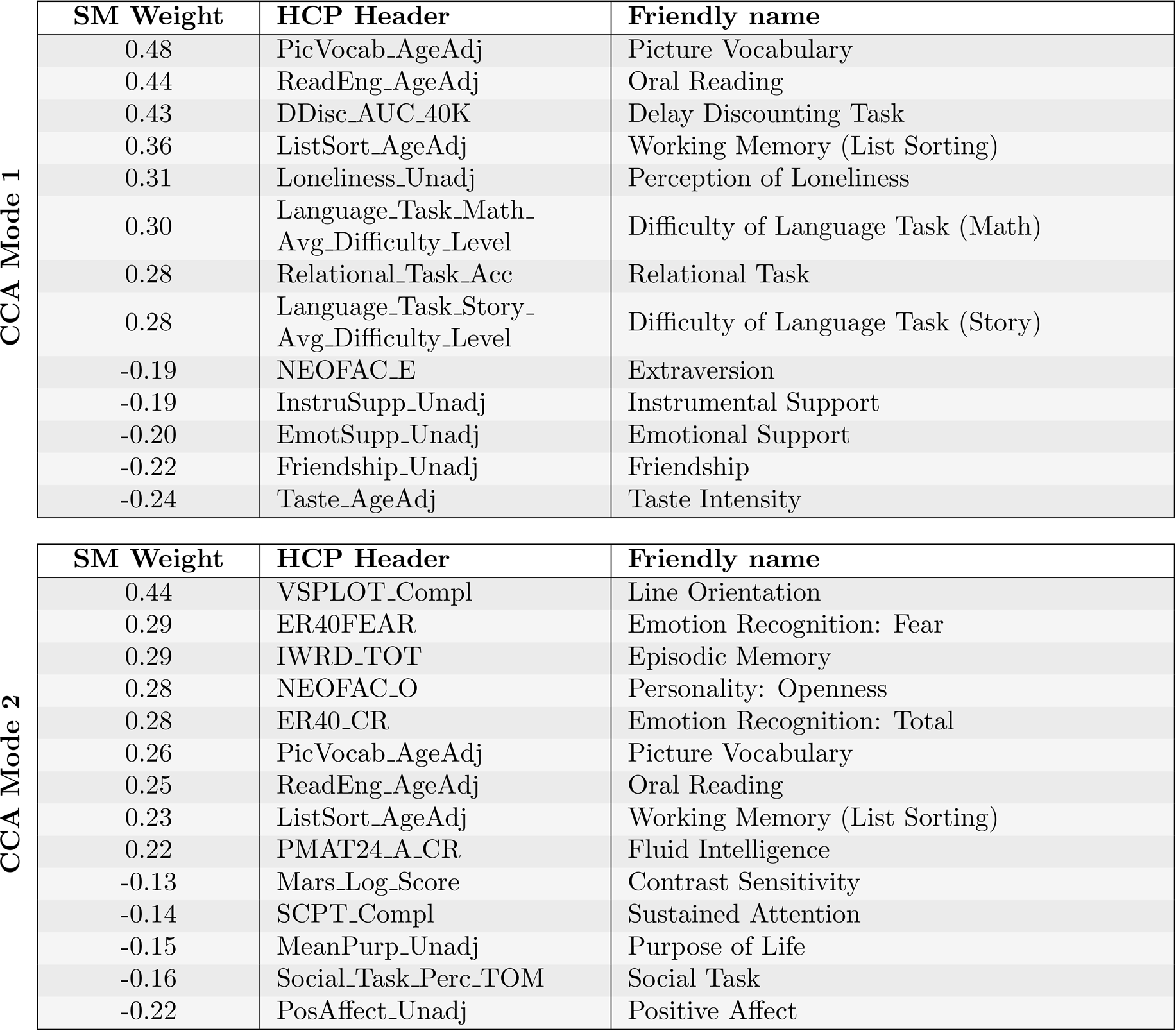
Fourth and fifth group-level dynamic modes of resting-state fMRI time series. The fourth mode is purely real, hence its period T is infinite, whereas the fifth mode is complex.

Figure S2 shows the second group-level dynamic modes for each of the five tasks considered in this work.

**Figure S2:**
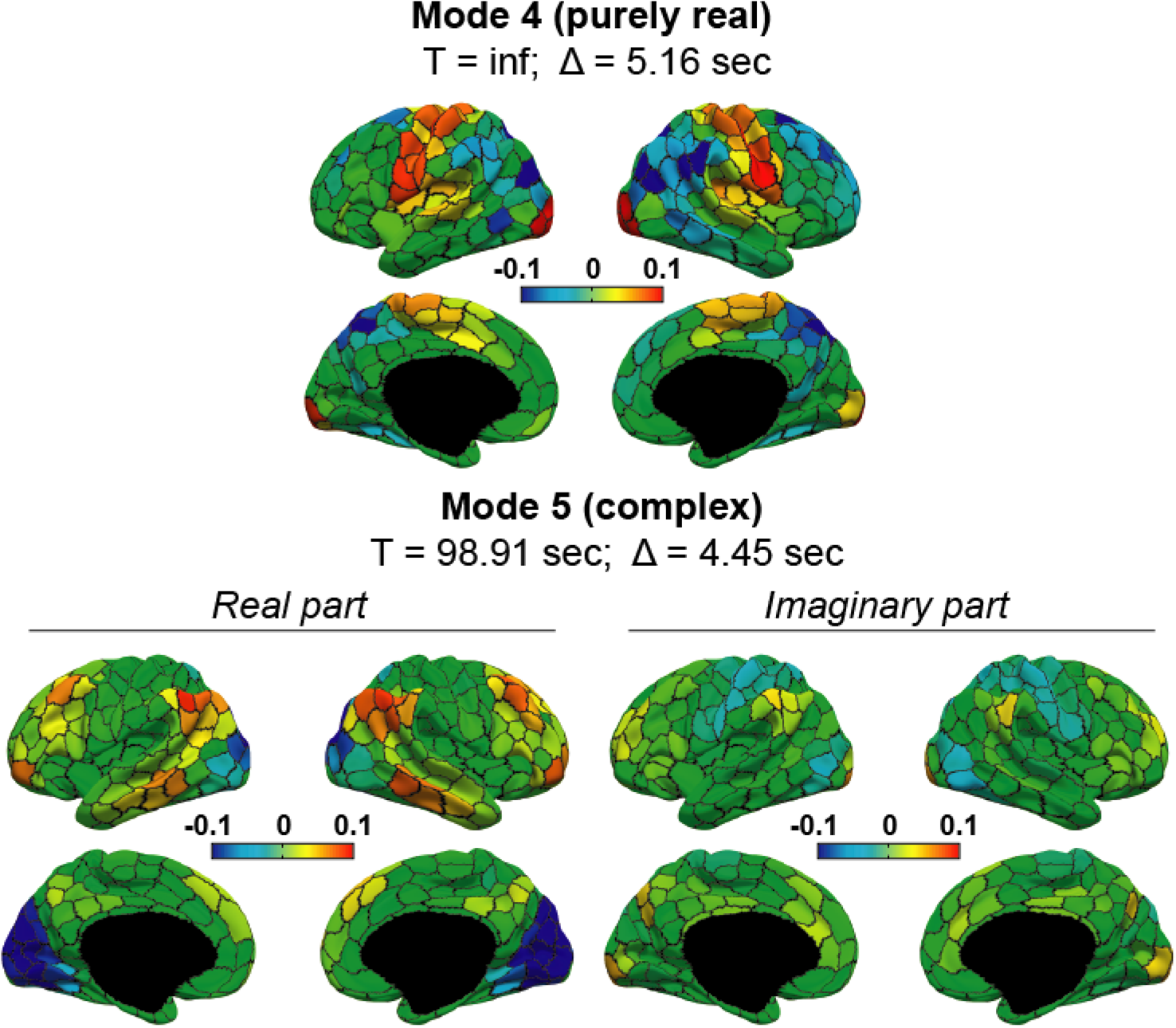
Second group-level dynamic modes for each of the five tasks considered in the present work and consisting in moving the left and right hands, the left and right feet, and the tongue.

### Impact of the length of the time series on detection of task

We test whether finding task-related activations in the imaginary parts of the dynamic modes is not due to a lucky choice of the fMRI time series’ 6-seconds subsections selected to compute the modes. To this end, we compute the first dynamic mode in each task using shifted subsections of different lengths of the corresponding time series. Figure S3 shows the first dynamic modes in the five task-conditions using the last 3-seconds subsections of the 6-second sections used originally. Activations directly related to the tasks and classically found in the corresponding activation maps (Yeo et al., 2011; Barch et al., 2013) are consistently encoded in the imaginary parts of the modes.

**Figure S3:**
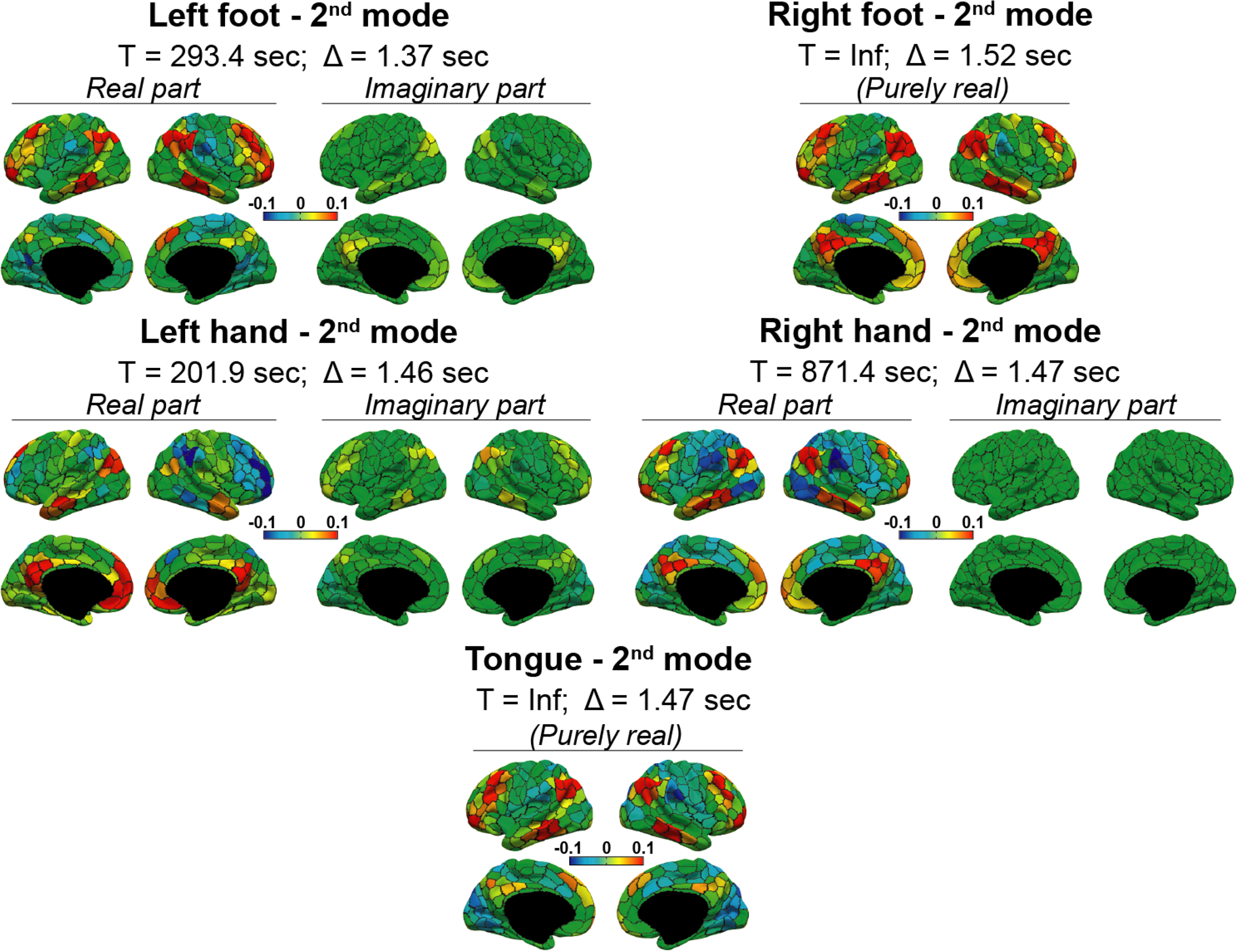
The task-related activation is also encoded in the imaginary part of the modes when only the last three seconds of the corresponding time series are considered.

We get similar results when using 4-seconds or 5-seconds subsections with different starting points within the 6-seconds windows.

### Distribution of damping times of resting-state and task dynamic modes

The distribution of the damping times of the first 100 dynamic modes in task and rest are presented in Figure S4. These distributions have similar shapes and suggest that the most important modes are found within the first 10–20 modes. Note that the vertical axes have different scales in rest and in task.

**Figure S4:**
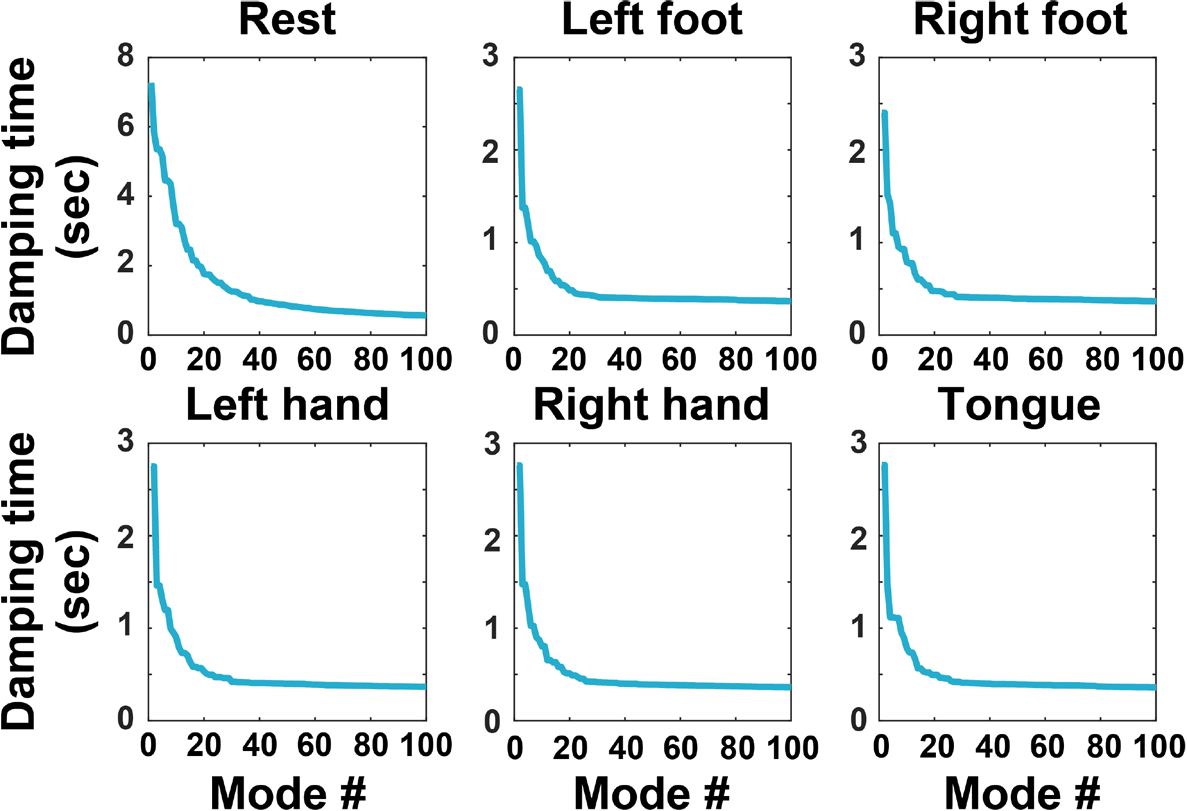
Damping times of the first 100 dynamic modes, ordered by decreasing values of damping times, in resting-state and in the five task conditions considered in this work.

Figure S5 shows the values of the frequencies associated to the first 100 dynamic modes.

**Figure S5:**
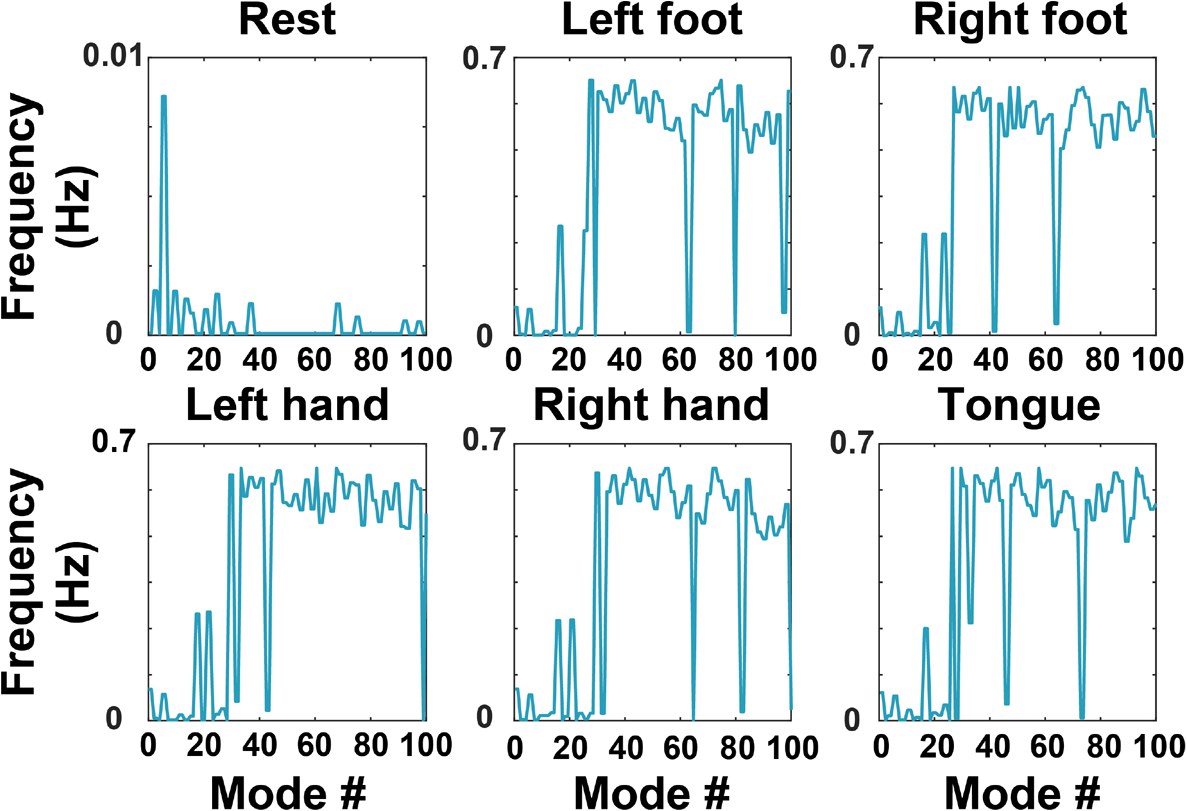
Frequencies associated to the first 100 dynamic modes, ordered by decreasing values of associated damping times, in resting-state and in the five task conditions considered in this work.

